# Contemporary Victoria and Yamagata Influenza B Viruses Elicit Lineage-Specific Differences in Innate Immunity

**DOI:** 10.64898/2026.09.17.752522

**Authors:** Caroline K. Page, Victoria Zyulina, Wuji Zhang, Ginger Geiger, Vrushali Dhamapurkar, Dabeiba Bernal- Rubio, David Guerrero, Sean D. Ray, Justin D. Shepard, Ana Fernandez-Sesma, Katherine Kedzierska, Stephen M. Tompkins

**Affiliations:** Center for Vaccines and Immunology, University of Georgia, Athens, Georgia 30602, USA; Center for Influenza Disease and Emergence Response (CIDER), University of Georgia, Athens, Georgia 30602, USA; Department of Infectious Diseases, University of Georgia, Athens, Georgia 30602, USA; Department of Microbiology Icahn School of Medicine at Mount Sinai. One Gustave L. Levy Place, Box1124, New York, NY 10129 USA; The Graduate School of Biomedical Sciences at Icahn School of Medicine. One Gustave L. Levy Place, New York, NY 10129 USA; Department of Microbiology and Immunology, University of Melbourne, Peter Doherty Institute for Infection and Immunity, Melbourne 3000 Victoria, Australia; HKU-Pasteur Research Pole, School of Public Health, LKS Faculty of Medicine, The University of Hong Kong, Hong Kong SAR, China

## Abstract

The two influenza B virus (FLUBV) lineages, Victoria and Yamagata, have continued to diverge since their separation in the 1970s, resulting in distinct antigenic characteristics and differences in immunity and cross-protection. Human epidemiological observations and experimental studies have identified lineage-specific differences in adaptive immune responses elicited by contemporary viruses, suggesting that innate immune signaling may contribute to these divergent outcomes. To understand in-depth immunity to contemporary influenza B viruses, we compared viral replication, cytokine production, gene expression, and cellular tropism following infection with representative Victoria- and Yamagata-lineage viruses. We found that, despite broadly similar cytokine profiles in the ferret upper respiratory tract, the two lineages exhibited distinct innate immune kinetics. Yamagata viruses induced rapid early expression of antiviral and inflammatory genes, including SOCS1 and multiple interferon-stimulated genes, whereas Victoria viruses displayed delayed innate immune activation accompanied by greater viral replication. Consistent with the ferret data, Yamagata viruses also induced elevated SOCS1 expression in human PBMCs early after infection. In addition, Yamagata viruses exhibited broader cellular tropism, infecting a wider range of immune cell populations than Victoria viruses. Together, these findings demonstrate that contemporary influenza B virus lineages distinctly engage the host immune system and provide new insight into how lineage-specific innate immune responses may contribute to differences in immunity, cross-protection, and the divergent evolutionary trajectories of Victoria and Yamagata viruses

**Author Summary:** Influenza B viruses cause substantial seasonal disease, yet the biological differences between the Victoria and Yamagata lineages that may contribute to their distinct epidemiology and immune responses remain incompletely understood. Recent studies comparing lineage-specific immunity have focused primarily on adaptive responses, leaving differences in early host–virus interactions comparatively unexplored. Here, we show that representative Victoria and Yamagata viruses differ during the earliest stages of infection. Using ferret and human immune cell models, we found differences in how the viruses trigger antiviral defenses, how efficiently they replicate, and which immune cells they infect. These findings suggest that early interactions between influenza B viruses and the immune system may help shape the distinct immunity observed between the lineages. Understanding these differences provides new insight into influenza B virus biology and may help inform our understanding of viral evolution, immunity, and future influenza vaccine development.

## Introduction

Influenza B viruses (FLUBVs) co-circulate with influenza A viruses (FLUAVs), contributing substantially to seasonal epidemics and disproportionately affecting vulnerable populations. Historically, FLUBVs have been understudied due to their lack of pandemic potential. However, they account for approximately 23% of influenza-related illnesses annually (1), and are increasingly recognized as a significant cause of morbidity and mortality, particularly in children (2-4). The earliest FLUBV was isolated in the 1940s (5) and by the 1980s, FLUBVs had diverged into two antigenically distinct lineages, Victoria and Yamagata (6) These lineages co-circulated globally until 2020, when Yamagata was last isolated from a human infection (7). Historically, FLUBVs underwent frequent reassortment events (8) including a complete neuraminidase gene segment swap in the early 2000s (9). However, since 2015, these lineages have followed distinct evolutionary trajectories, culminating in the apparent disappearance of Yamagata from human circulation. These evolutionary changes have likely influenced virus-host interactions and immune responses, yet the mechanisms underlying lineage-specific immunity remain poorly understood (10, 11).

The epidemiology of FLUBVs has changed substantially since the COVID-19 pandemic. With no confirmed detection of Yamagata since March 2020, the World Health Organization has recommended the removal of the Yamagata component from seasonal influenza vaccines and the transition from quadrivalent to trivalent formulations(12). Unlike FLUAVs, which are maintained in diverse animal reservoirs, FLUBVs are largely restricted to humans and depend almost exclusively on human-to-human transmission for persistence. Consequently, their evolutionary success is closely tied to their ability to interact with and evade the host immune responses. Defining how the Victoria and Yamagata lineages differentially engage innate and adaptive immune responses is therefore critical for understanding viral fitness, transmission, and pathogenesis, and future vaccine design.

We have previously shown in mice and ferrets that there are differences in cross-protection between the FLUBV lineages, attributed to the adaptive humoral immune responses (13). These data corroborate human epidemiological observations which found that FLUAV types and Victoria viruses are effective at providing protection against reinfection in subsequent seasons whereas Yamagata viruses are insufficient in conferring the same level of immunity (14). While the adaptive immune response is key in preventing infection and reducing disease severity, the innate immune response plays a critical role in shaping adaptive immunity, influencing the quality and longevity of protective responses through cytokine signaling and antigen presentation.

In depth characterization of innate immunity contributes to understanding disease severity, pathogenicity, and protection from future infections. Studies comparing FLUAV and FLUBV infections have found that FLUBVs cause milder pathogenesis, weaker inflammatory responses, and reduced neutralizing antibody titers in ferrets following infection (15-17). FLUBV infection also modulates gene expression differently from FLUAV by inducing faster IFN regulatory factor 3 (IRF3) and IFN-λ1 expression suggesting differential viral entry (18, 19). While these studies have highlighted key immunological differences between influenza types, less is known about the innate immunological differences between the two influenza B lineages. Furthermore, the divergence of contemporary isolates may reflect changes in innate immune signaling, including variations in cytokine and chemokine production, as well as shifts in gene regulation, which could significantly impact disease dynamics.

Ferrets are widely recognized as an excellent model for studying influenza due to their close resemblance to humans in both respiratory physiology and immune response (20, 21). Unlike many other small animal models, ferrets are naturally susceptible to infection with human influenza viruses without prior adaptation, allowing for direct study of viral replication, transmission, and pathogenesis (22). Ferrets also exhibit clinical signs of influenza infection, such as fever, lethargy, and sneezing, which parallel symptoms in humans, providing a robust system for evaluating disease severity. However, a significant limitation of using ferrets as a model is the lack of species-specific immunological reagents and assays, which restricts the ability to comprehensively analyze immune responses at a cellular and molecular level. Despite these challenges, recent advances in ferret-immune technology have enabled a deeper immunological assessment of influenza infection, paving the way for more detailed investigations into host-pathogen interactions.

Given the differences in adaptive immunity and cross-protection, we sought to investigate how distinct innate immune responses between contemporary lineage viruses might contribute to these observed variations in protection and disease dynamics. Our findings demonstrate that the two lineages exhibit distinct cytokine and gene expression kinetics, with Yamagata lineage viruses upregulating key innate antiviral genes early after infection in ferrets. Complementary experiments using human peripheral blood mononuclear cells (PBMCs) and monocyte-derived dendritic cells (moDCs) corroborated ferret data by revealing upregulation of certain genes, such as SOCS1, further supporting lineage-specific immune modulation, and highlighted differential infection tropism between the viruses. These data underscore lineage-specific differences in innate immune responses, which may provide insights into the long-term infection dynamics and the disappearance of B/Yamagata viruses from human circulation.

## Results

### Victoria-lineage Virus has Overall Greater Viral Replication in Ferret’s Upper Respiratory Tract

To assess differences in viral replication of a contemporary Victoria and Yamagata lineage virus, ferrets were inoculated intranasally with either B/Washington/02/2019 (Victoria) or B/Oklahoma/10/2018 (Yamagata) at 10^6^ PFU (plaque-forming units). Nasal wash samples were collected on days 1, 3, 5 and 7 post-infection to assess viral shedding in the upper respiratory tract (URT) and animals were monitored for clinical signs, including weight loss and temperature changes. No significant differences in clinical signs were observed between the two groups, as neither infection caused substantial weight loss **(Fig. 1A)** or changes in temperature **(Fig. 1B)**. On day 3 post-infection, ferrets infected with Yamagata showed a slight decrease in weight, which returned to baseline or surpassed baseline by day 5. In contrast, Victoria-infected ferrets exhibited a spike in temperature on day 5, which subsequently decreased to near baseline levels by day 7.

**Figure 1.**
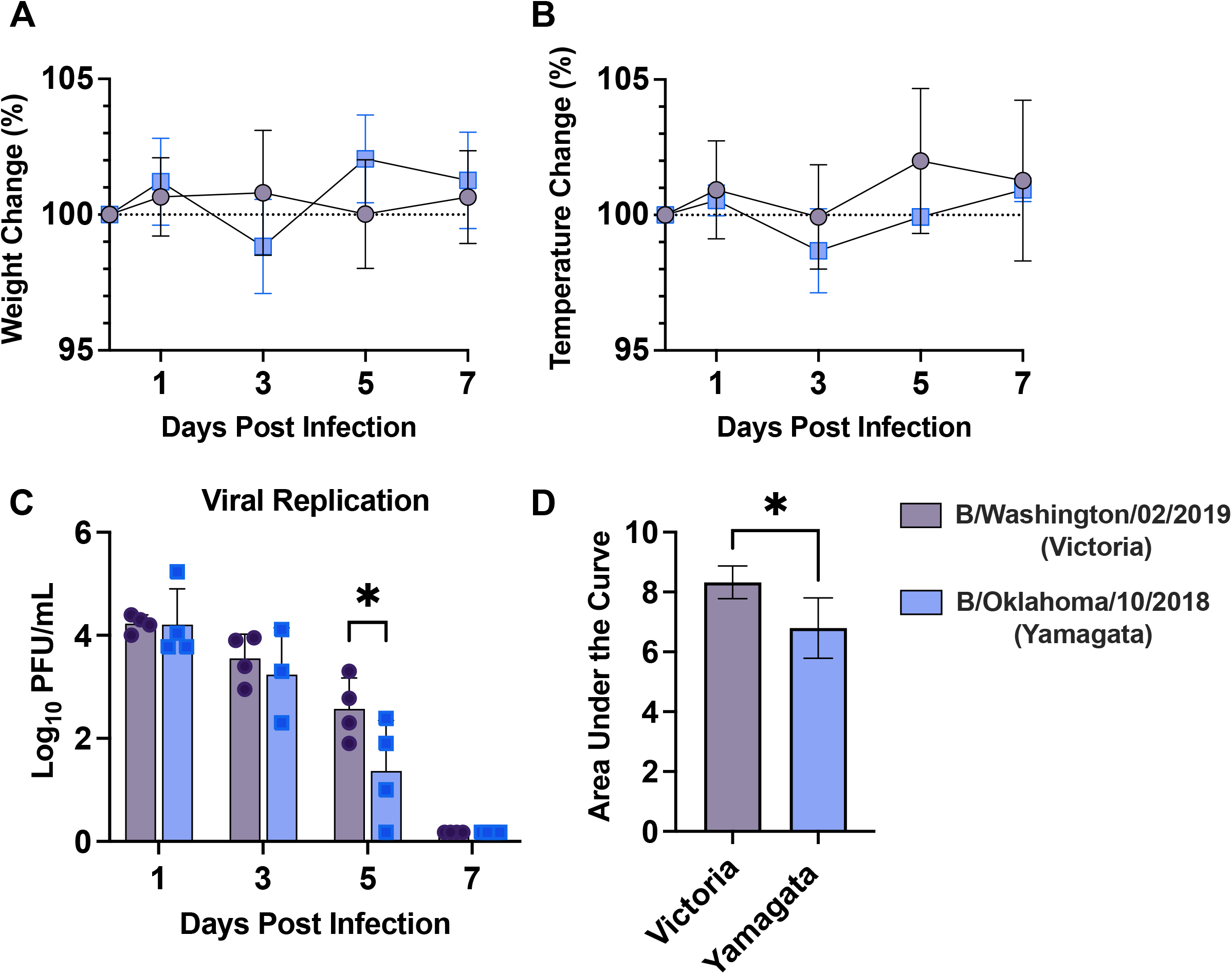
Victoria-lineage Virus has Greater Replication in Ferrets. A. Weight loss and B. Temperature changes in ferrets following infection with 10^6^ PFU of B/Washington/02/2019 (purple) or B/Oklahoma/10/2018 (blue). C. Viral replication of FLUBV infection in the nasal wash samples of ferrets on days 1, 3, 5, and 7 post-infection, measured by plaque assay. D. Total viral replication, measured as area under the curve of C, following FLUBV infection. Statistical significance was determined using a two-way ANOVA for viral replication and a t-test for area under the curve analysis.

Viral replication in the URT were similar for both infections, with peak viral shedding occurring on day 1 and clearance by day 7. The Victoria-lineage virus displayed significantly higher viral replication on day 5 post-infection compared to the Yamagata-lineage virus **(Fig. 1C)**. The area under the curve analysis (AUC) revealed total viral replication was significantly higher in the Victoria-infected ferrets than in the Yamagata-infected ferrets indicating that Victoria exhibited better overall replication than Yamagata in ferret models **(Fig. 1D)**.

### Yamagata Lineage Virus Showed Delayed Chemokine and Cytokine Responses in Ferret URT

Cytokine protein expression in the URT was measured using a multiplex assay on ferret nasal wash samples collected on days 1, 3, 5, and 7 post-infection **(Fig. 2 A-H)**. The Victoria-lineage infection demonstrated peak expression of most cytokines on day 3, including IL-8, IL-12p70, IL-2, IL-6, CXCL10 (IP-10), CCL2 (MCP-1), and TNF-α **(Fig. 2 B-H)**. In contrast, the Yamagata-lineage infection showed peak expression of cytokines primarily on day 5, including IL-8, IL-12p70, IL-6, CXCL10, CCL2, and TNF-α **(Fig. B, C and E-H)**. Expression of IFN-γ peaked on day 5 for both the Victoria and Yamagata infection **(Fig. 2A)**. By day 7, expression levels induced from both infections were minimal, indicating resolution or reduction of cytokine production. The altering kinetics between infections is exemplified by the TNF-α expression **(Fig. 2H)**. Overall TNF-α concentration levels were comparable but the peak occurred on day 3 for Victoria infection and day 5 for Yamagata infection.

**Figure 2.**
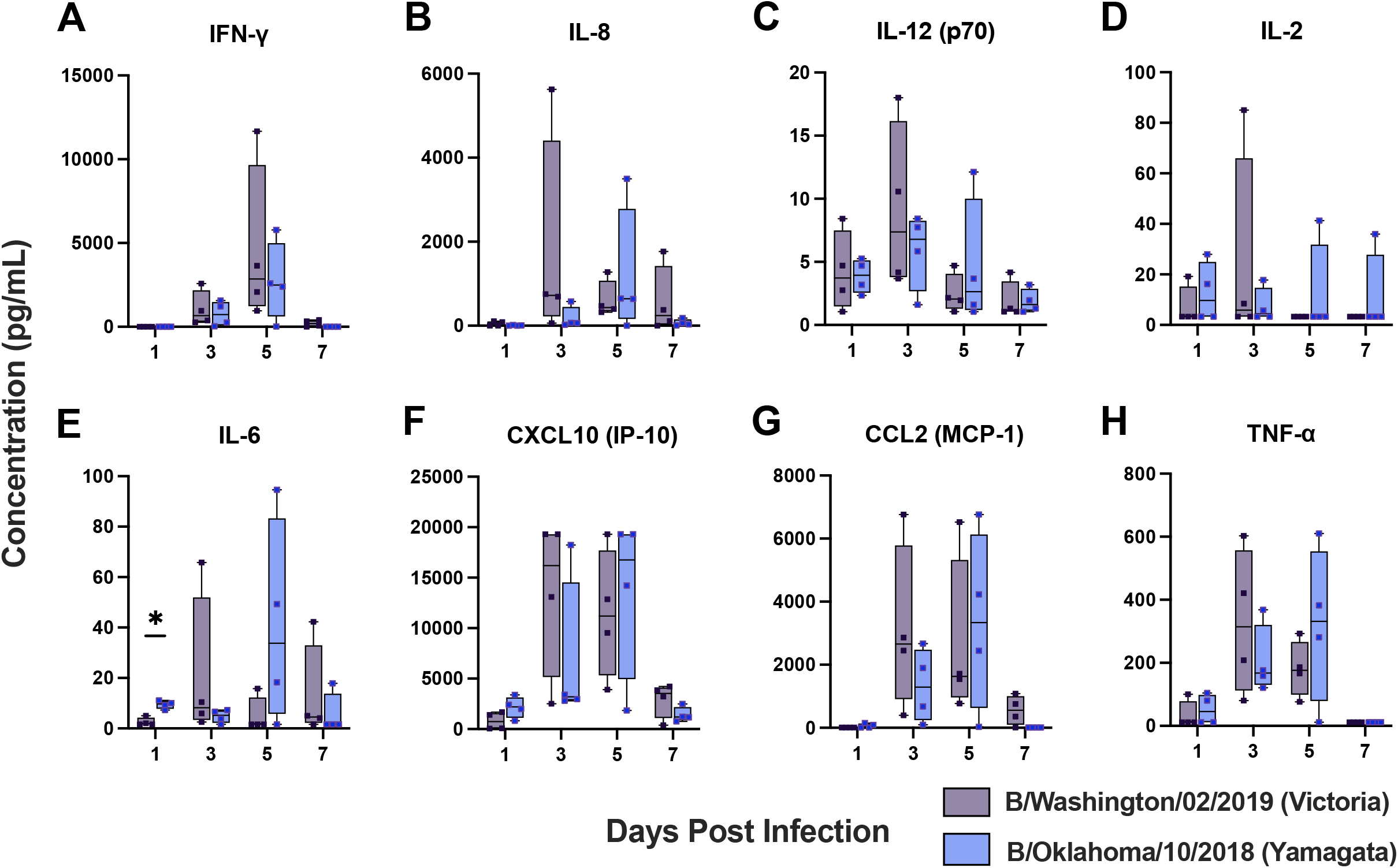
Cytokine and Chemokine Levels in Ferrets Following FLUBV Infection. Cytokine and chemokine levels were measured in the nasal wash samples of ferrets on days 1, 3, 5, and 7 post-infection using a Luminex multiplex assay. B/Washington/02/2019 represents the Victoria virus (Purple) and B/Oklahoma/10/2018 represents the Yamagata virus (Blue). Values for chemokines and cytokines are graphed as concentrations in pg/mL. Statistical analysis was performed using a two-way ANOVA.

Notably, overall IFN-γ expression levels trended higher in the Victoria-infected ferrets compared to Yamagata-infected ferrets although not significantly different **(Fig.2A)**. Similarly, IL-12p70 and IL-2, cytokines associated with Th1 responses, trended higher in Victoria infections compared to Yamagata infections on their respective peak days **(Fig. 2C and E)**.

### Transcriptional Regulation of Innate Immune Responses During FLUBV Infection in ferrets

Correlations between protein levels and gene expression are not always consistent **(23-25)**. To better understand the transcriptional regulation and potential upstream signaling pathways contributing to the differential responses elicited by each lineage virus, we analyzed gene expression profiles in the URT during infection using a QuantiGene Plex Gene Expression Assay. A custom panel of 50 genes was designed to comprehensively assess the innate immune response to infection with contemporary FLUBV lineage viruses **(Fig. 3)**. The custom panel was selected to include key components of the innate immune response, focusing on antiviral signaling pathways, cytokine activity, and genes previously implicated in FLUBV infections.

**Figure 3.**
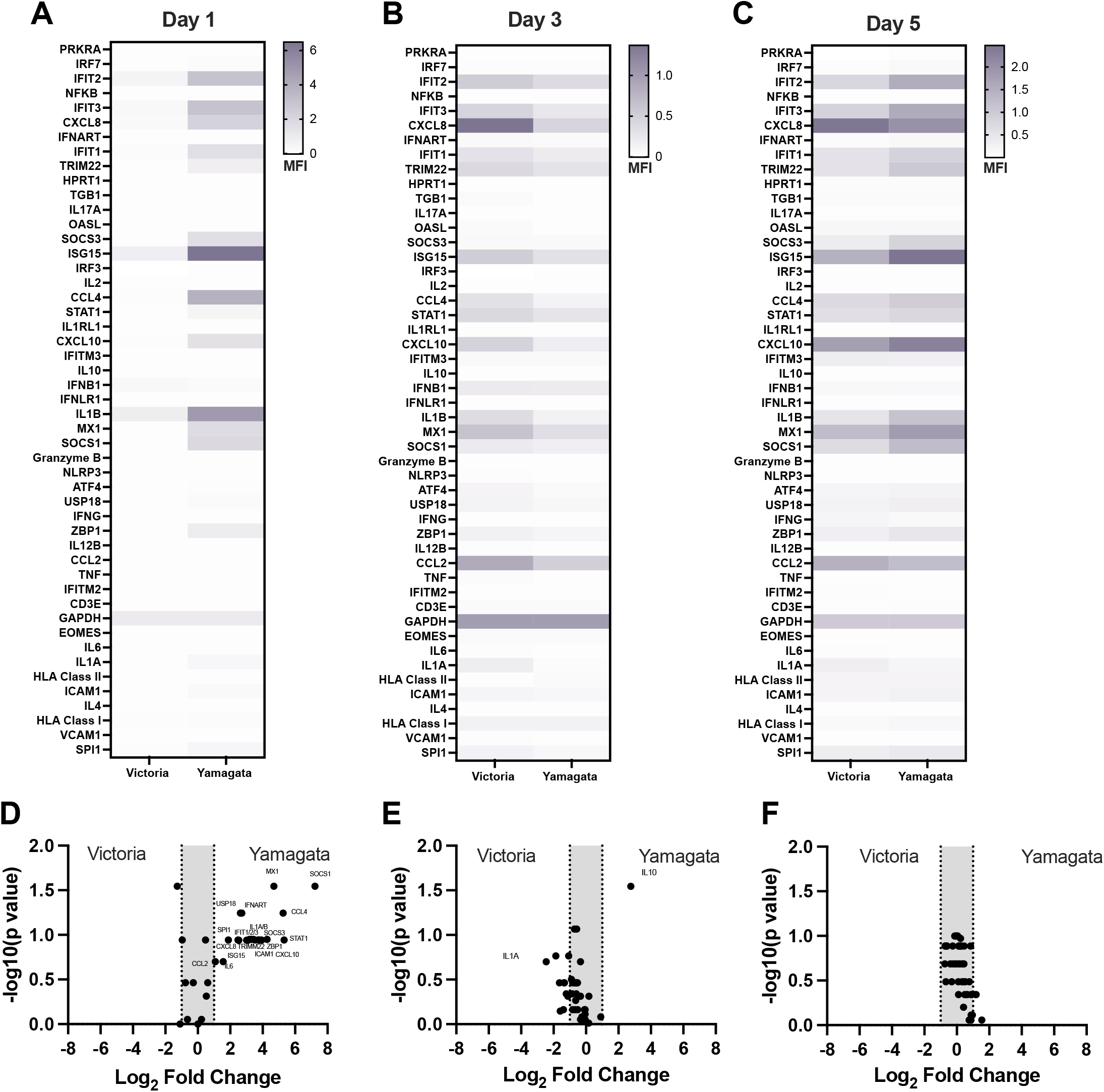
Gene Expression between Victoria and Yamagata Lineages. Following Infection in Ferrets. A-C. Mean fluorescent intensity (MFI) of each gene (y-axis) measured from RNA isolated from nasal wash samples on days 1, 3 and 5 post-infection with a Victoria (B/Washington/02/2019) or Yamagata (B/Oklahoma/10/2018) virus (x-axis). The signal was normalized to housekeeping gene, GAPDH, for each day, respectively. Darker purple represents stronger signals, while white represents weaker signals. D-E Volcano plots of differential gene expression of a Victoria infection compared to a Yamagata infection on days 1, 3, and 5 post-infection. P-values are on the y-axis while the log_2_ fold changed of each gene (Victoria vs Yamagata) is represented on the x-axis. Significantly differential expressed genes are labeled on the graph.

In contrast to the low or undetectable cytokine protein levels on day 1 post infection, the gene expression levels revealed significant differences between the viruses. An increase in the expression of IFIT2, IFIT3, CXCL8, IFIT1, TRIM22, SOCS3, ISG15, CCL4, CXCL10, IL1B, MX1, and SOCS1 was observed day 1 post infection in the upper respiratory tract of Yamagata-infected ferrets compared to Victoria-infected ferrets **(Fig. 3A)**. By day 3, these differences in gene expression were reduced as the Victoria infection began to drive increased expression of key antiviral genes, including IFIT2, IFIT3, CXCL8, TRIM22, ISG15, MX1, and CCL2, among others **(Fig. 3B)**. Notably, some of these genes showed expression levels surpassing those induced by the Yamagata infection. By day 5, gene expression levels between the two infections were comparable, with consistently high expression of certain antiviral genes, including IFIT2, IFIT3, CXCL8, ISG15, CXCL10, MX1, and CCL2 **(Fig. 3C)**.

When assessing the differentially regulated genes between the infections on each day, we observed a trend consistent with the normalized gene expression analysis: day 1 post-infection exhibited the highest number of differentially regulated genes, with these differences diminishing or disappearing by days 3 and 5 **(Fig. 3D-F)**. On day 1, Yamagata infection significantly upregulated several key innate immune regulatory genes, including IFIT1, IFIT2, IFIT3, ISG15, CXCL8, SOCS1, SOCS2, TRIM22, CXCL10, USP18, STAT1, CCL4, and MX1. Among these, SOCS1 was the most significantly upregulated gene when comparing the two infections.

### Gene Expression and Cellular Tropism of Victoria and Yamagata Lineage Influenza B Viruses in Human PBMCs

In ferrets, we identified multiple genes that were differentially regulated early after infection with Victoria and Yamagata lineage viruses at the site of infection in the URT. To translate these findings to human systems, we infected human peripheral blood mononuclear cells (PBMC) with a representative Victoria (B/Washington/02/2019) and Yamagata (B/Sydney/701/2018) lineage virus and analyzed gene expression for previously identified key genes: IFIT1, IFIT3, IFN-y, SOCS1, MX1, ISG15, TNF and STAT1. PBMCs from five unique donors were infected with a multiplicity of infection (MOI) of 4 and samples were collected for RT-qPCR analysis at 8- and 22-hours post-infection **(Fig. 4)**. Notably, SOCS1 stood out as a gene which was statistically significantly increased in Yamagata-infected PBMCs compared to Victoria-infected PBMCs at 8 hours post-infection **(Fig. 4A-B**). By 22 hours post infection, like our later timepoint observations in the ferret data, we observe significant increases expression of IFNy and ISG15 for the Victoria lineage virus compared to the Yamagata virus **(Fig. 4C-D)**.

**Figure 4.**
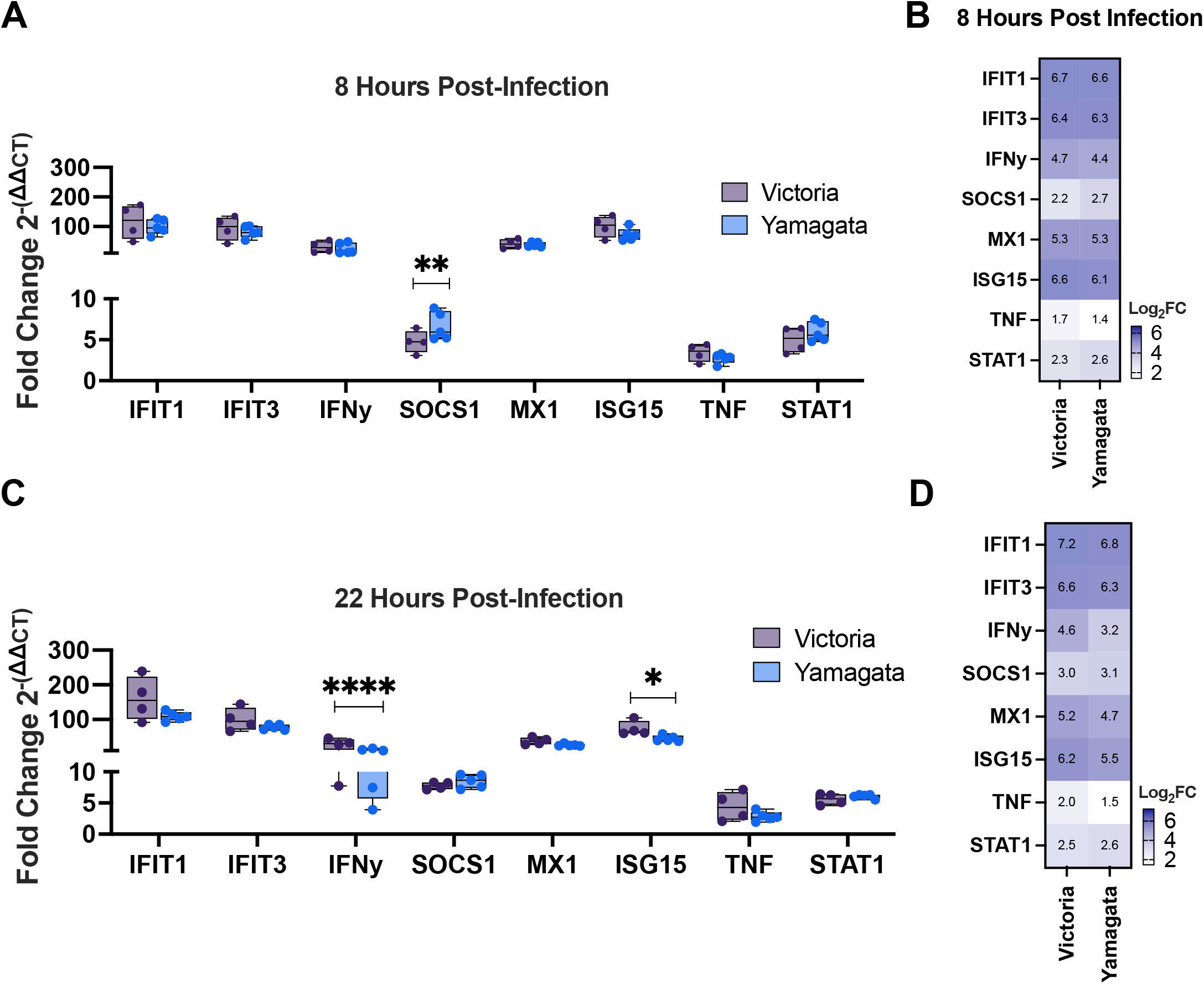
Gene Regulation in Human PBMCs Following Influenza B Virus Infection. Bar graph and heatmap representing qPCR analysis of differentially expressed genes in human PBMCs following infection with contemporary influenza B viruses at (A-B) 8- or (C-D) 22-hours post-infection. Infection groups include Victoria-lineage (B/Washington/02/2019; purple) and Yamagata-lineage (B/Sydney/701/2018; blue) at an MOI of 4. Gene expression was normalized to the geometric mean of GAPDH and 18S and is presented as Fold Change 2^-(ΔΔCT)^ and Log_2_FC for clarity. Statistical significance was determined using one-way ANOVA (*p* < 0.05). Error bars represent the minimum, maximum, and interquartile range of 5 biological replicates.

To assess possible differences in the cellular tropism of each virus, PBMCs infected with an MOI of 4 of each virus were collected 22-hours post infection and nucleoprotein (NP) positive cells were determined by flow cytometry. Cells were gated using monocyte, natural killer (NK) cell, B cell, γδ T cell, CD4+ T cell, and CD8+ T cell-markers, within the nucleoprotein (NP)-positive population **(Fig. 5A and B)**. All viruses, Victoria, Yamagata, and the control FLUAV (A/California/07/2009 (pdmH1N1) with PR8 backbone), could efficiently infect monocytes, with Victoria-infected PBMCs showing significantly more NP+ monocytes than Yamagata **(Fig. 6C)**. Notably, the Yamagata lineage virus uniquely demonstrated the ability to efficiently infect a broad range of immune cells, including B cells, T cells, and NK cells. At 22 hours post-infection, Yamagata-infected PBMCs exhibited significantly increased frequencies of NP+ NK cells, γδ T cells, B cells, CD8+ T cells, and CD4+ T cells compared to the Victoria lineage and the FLUAV control virus **(Fig. 5D-H)**. A t-SNE analysis of the flow data, which is a nonlinear dimensionality reduction technique used to help visualize high dimensional datasets, independently clustered the immune cell groups and showed the broad cellular tropism displayed by the Yamagata virus compared to the Victoria and FLUAV control **(Fig. 5I-L)**

**Figure 5.**
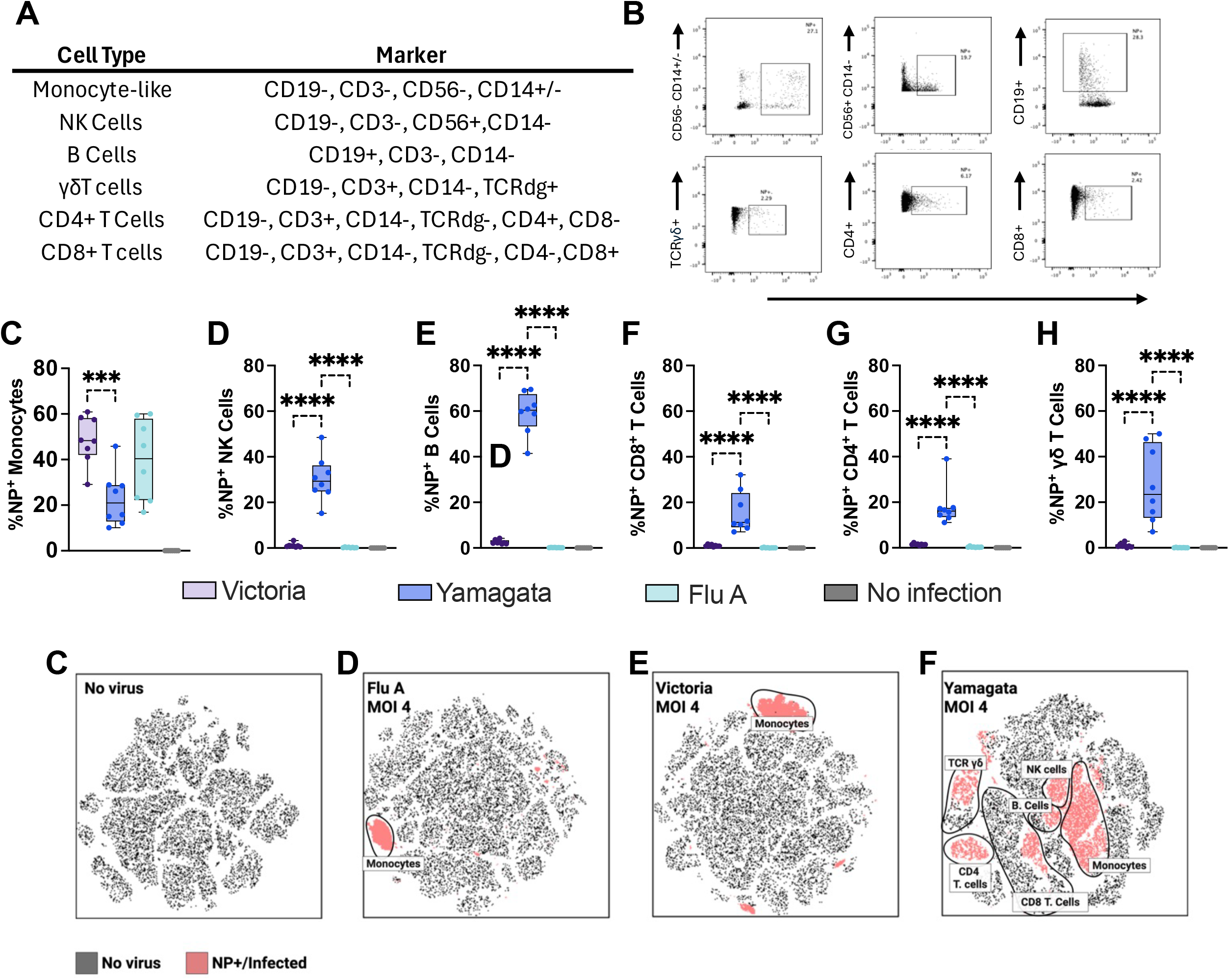
Differential Infection of Immune Cell Types Following FLUBV Infection. A. Table summarizing the flow cytometry strategy used to distinguish immune cell types and the markers used for their identification. B. Representative dot plots for NP+ populations from each cell type. C-H. Bar graphs showing the percentage of NP+ cells from each cell type following an infection with A/California/07/2009 (Flu A; Light Blue), B/Sydney/701/2018 (Yamagata; Dark Blue), B/Washington/02/2019 (Victoria; Purple), and no virus (Grey) at 8- and 22-hours post-infection with an MOI of 1 or 4. I-L. tSNE plots visualizing clustering of infected (light pink) and uninfected (black) cell populations.

**Figure 6.**
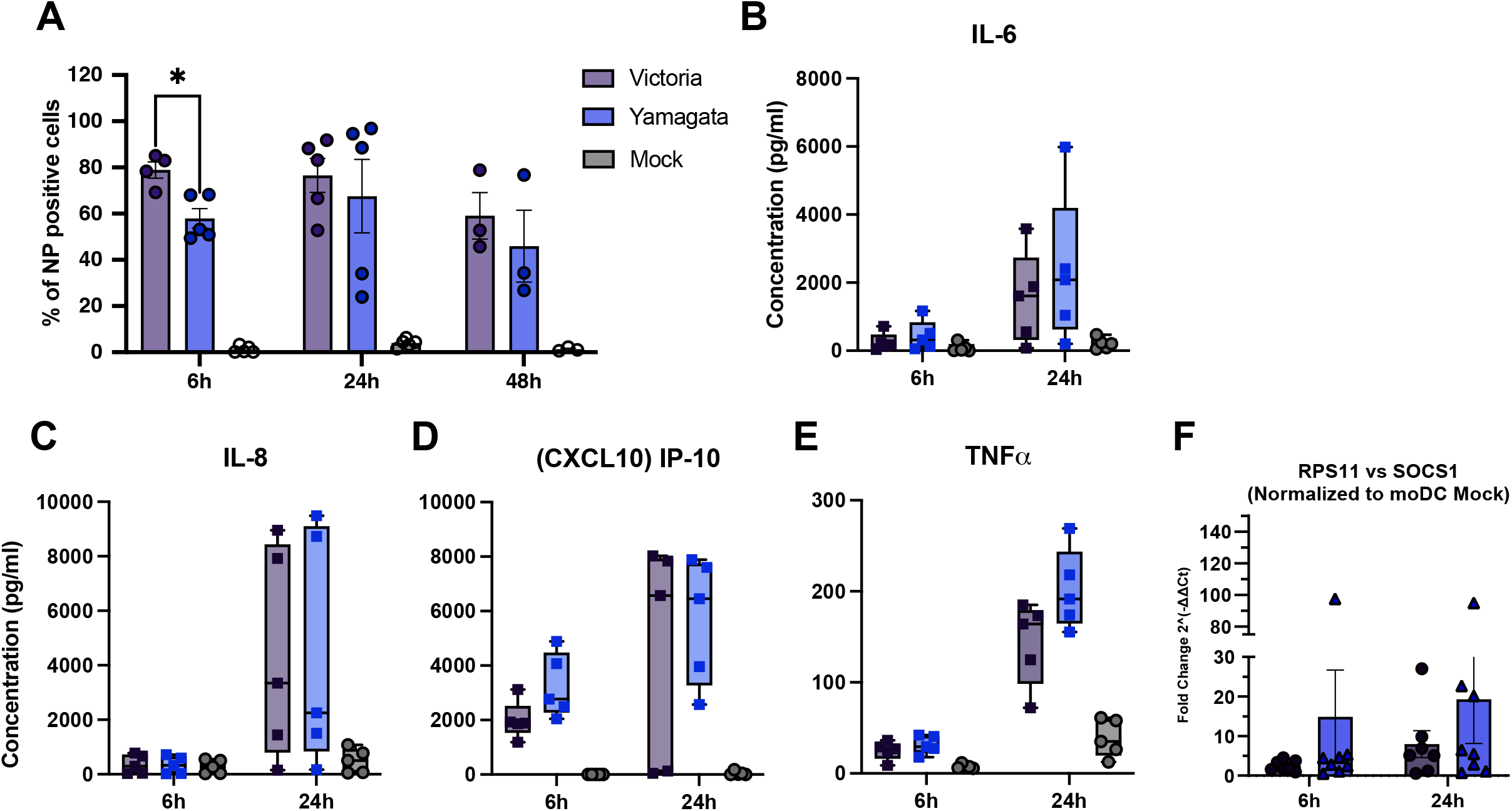
SOCS1 expression in monocytic derived dendritic cells. A. Frequency of NP-positive MoDCs following infection with Victoria- or Yamagata-lineage influenza B viruses (MOI = 1) at the indicated time points. B–E. Concentrations of IL-6, IL-8, CXCL10, and TNF-α in culture supernatants measured by multiplex ELISA. F. Relative SOCS1 expression measured by RT-qPCR and normalized to RPS11 and mock-infected controls. Data represent five independent donors. Symbols represent individual donors and bars indicate mean ± SEM. Statistical significance was determined by two-way ANOVA with multiple comparisons.

### SOCS1 expression in monocytic derived dendritic cells

Monocytic-derived dendritic cells (MoDCs) are antigen-presenting cells differentiated from circulating monocytes following infection or stimulation (26). They play an important role in bridging the innate and adaptive immune responses and are known to express high levels of SOCS1 (27). Since our human PMBC infection data showed that monocytes were the predominant cell type infected by both FLUAV and FLUBVs **(Figure 5)**, we next sought to further characterize the response of MoDCs to FLUBV infection. To address this, we infected MoDCs and compared susceptibility to infection, cytokine production, and SOCS1 expression between the FLUBV lineages. MoDCs from five human donors were infected with either Victoria or Yamagata lineage virus at an MOI of 1, and cells were collected at 6, 24 and 48 hours-post infection to quantify NP+ cells. The Victoria lineage virus resulted in significantly higher frequencies of NP+ cells at 6 hours post-infection compared to Yamagata, although this difference was no longer apparent at 24 and 48 hours **(Fig. 6A)**. Despite this difference in early infection, Mo-DCs produced comparable levels of IL-6, IL-8, CXCL10, and TNF-α following infection with either virus at all timepoints **(Fig. 6B-E)**. SOCS1 expression was assessed at 6- and 22-hours post-infection by RT-qPCR, normalized to the housekeeping gene ribosomal protein S11 (RPS11) and mock-infected cells. Although no significant differences in SOCS1 expression were observed between the two lineages, expression increased over time, with SOCS1 levels trending higher following Yamagata infection **(Fig.6F)**.

## Discussion

Contemporary influenza B virus lineages have diverged in ways that alter immunity and cross-protection between them, potentially contributing to the disappearance of the Yamagata lineage from human circulation. Continued animal and translational studies are therefore essential to better define the biological and immunological differences between these viruses. To address this, we compared host cytokine and gene expression responses following infection with contemporary Victoria- and Yamagata-lineage influenza B viruses in ferrets and human PBMCs. While all animal models have limitations, ferrets are widely regarded as one of the most relevant models for studying influenza disease and immunity because they recapitulate many aspects of human infection without requiring viral adaptation(20, 21). Limited studies have explored innate immune responses to influenza B virus infection in ferrets, largely due to the scarcity of ferret-specific immunological reagents. A 2016 study by Carolan et al. found no major differences in local immune responses in the upper and lower respiratory tracts of ferrets infected with A(H1N1)pdm09, A(H3N2), or a Yamagata-lineage influenza B virus (28). However, a Victoria-lineage virus was not included in that study, and the two influenza B virus lineages have diverged further in the years since. More recently, Rowe et al. used gene expression and cytokine profiling to compare influenza A and B virus infections in ferrets and identified subtype-specific differences; however, these analyses were performed in serum and did not evaluate local immune responses at the site of infection, the upper respiratory tract (17).

Although overall cytokine production was similar between the Victoria- and Yamagata-lineage infections, the kinetics of these responses differed substantially. The Victoria-lineage virus exhibited greater overall viral replication in the upper respiratory tract and induced higher peak expression of several pro-inflammatory cytokines, including IFN-γ, IL-12p70, and IL-2. However, cytokine responses peaked earlier during Victoria infection, occurring predominantly on day 3 post-infection, whereas peak cytokine expression following Yamagata infection was delayed until day 5. These findings suggest that the two lineages differ not necessarily in the magnitude of the inflammatory response they induce, but rather in the timing of innate immune activation.

Analysis of transcriptional responses further supported this observation. While cytokine protein measurements provide insight into biologically active immune mediators, transcript abundance does not always directly correlate with protein expression due to post-transcriptional regulation, differences in protein stability, and temporal delays between transcription and translation. Consistent with this, gene expression profiling revealed substantially greater differences between the two infections at early time points than were observed at the protein level. Yamagata infection induced robust early expression of antiviral and inflammatory genes, including IFIT1, IFIT2, IFIT3, ISG15, CXCL8, SOCS1, TRIM22, CXCL10, STAT1, and MX1, whereas Victoria infection induced comparatively limited early transcriptional responses despite exhibiting higher viral replication later in infection. By days 3 and 5, transcriptional responses between the two infections became increasingly similar. Together, these findings suggest that Yamagata viruses trigger a more rapid innate immune response, whereas Victoria viruses exhibit delayed innate immune activation that may contribute to enhanced viral replication in the upper respiratory tract.

Among the genes differentially expressed during early infection, SOCS1 was the most significantly upregulated transcript in Yamagata-infected ferrets. SOCS1 is a member of the suppressor of cytokine signaling family and functions as a negative regulator of cytokine signaling through inhibition of the JAK-STAT pathway, a central mediator of antiviral immunity (29, 30). Cellular pathway analysis further linked SOCS1 and SOCS3 to the PI3K complex, which plays important roles in both host antiviral responses and influenza virus replication (SI Figure 1) (31). Consistent with the ferret data, SOCS1 expression was also significantly elevated in Yamagata-infected human PBMCs early after infection, suggesting that enhanced SOCS1 induction may be a conserved feature of Yamagata-lineage virus infection across model systems. A similar trend toward increased SOCS1 expression following Yamagata infection was observed in MoDCs, although this difference did not reach statistical significance and considerable inter-donor variability was observed. This variability may reflect differences in the immune histories of human donors and highlights the complexity of translating lineage-specific responses observed in naïve animal models to human immune cells.

The biological consequences of this differential SOCS1 expression remain unclear. Because influenza viruses can modulate host signaling pathways to influence replication and host immune responses, it is possible that enhanced SOCS1 induction contributes to the distinct innate immune kinetics observed during Yamagata infection. In addition, although the NS1 proteins of the Victoria and Yamagata viruses used in this study are highly conserved, sequence differences within functional domains involved in host protein interactions may contribute to lineage-specific regulation of innate immune signaling. Future studies will be needed to determine whether viral factors such as NS1 directly influence SOCS1 expression and downstream antiviral responses.

Consistent with the differences observed in innate immune activation, the Victoria- and Yamagata-lineage viruses also differed in their cellular tropism. While both the H1N1 and Victoria-lineage viruses primarily infected monocytes, the Yamagata-lineage virus infected a broader range of immune cell populations, including NK cells, γδ T cells, B cells, CD8+ T cells, and CD4+ T cells. This expanded tropism may influence multiple arms of the immune response and could contribute to the distinct immune signatures observed following Yamagata infection. Because innate immune responses shape the magnitude and quality of adaptive immunity, these early differences in cellular targeting may have downstream consequences for lineage-specific immune responses.

Several limitations should be considered when interpreting these findings. The lineage-specific innate immune differences observed here likely represent only one component of the broader biological differences between Victoria and Yamagata viruses, as adaptive immune responses and other host-virus interactions were not evaluated. In addition, cellular tropism was inferred from intracellular NP staining and therefore may not exclusively reflect productive infection. Finally, these experiments were conducted using a single representative virus from each lineage; therefore, the observed differences may reflect strain-specific characteristics and should not be assumed to generalize across all Victoria- and Yamagata-lineage viruses.

In conclusion, our findings reveal clear lineage-specific differences in the innate immune responses elicited by contemporary Victoria- and Yamagata-lineage influenza B viruses. Although both viruses induced broadly similar cytokine profiles, Yamagata infection was characterized by rapid activation of antiviral transcriptional programs, including early induction of SOCS1 and interferon-stimulated genes, whereas Victoria infection exhibited delayed innate immune activation accompanied by greater viral replication in the upper respiratory tract. In addition, the two lineages differed in their cellular tropism, with Yamagata viruses infecting a broader range of immune cell populations than Victoria viruses. Together, these findings demonstrate that contemporary influenza B virus lineages engage the host immune system through distinct mechanisms and suggest that differences in early innate immune signaling may contribute to the divergent patterns of immunity and cross-protection previously observed between Victoria and Yamagata viruses. As the two lineages have followed markedly different evolutionary and epidemiological trajectories, culminating in the apparent disappearance of Yamagata viruses from human circulation, these results provide additional evidence that contemporary influenza B viruses are not immunologically interchangeable and underscore the importance of understanding lineage-specific virus–host interactions.

## Methods

### Viruses

Influenza viruses B/Washington/02/2019 (FR-1709) and B/Oklahoma/10/2018 (FR-1660) used for ferret infections in this study were obtained from the International Reagent Resource (IRR) and propagated in 9-to-11-day-old specific pathogen free embryonated chicken eggs according to established procedures (32). Viruses were sequenced to determine the egg passaging had minimal impact on viral adaptations. For human PBMC infections, A/California/07/2009 (PR8 backbone), B/Washington/2019, and B/Sydney/701/2018 were propagated in-house (Melbourne, Australia). For viruses propagated in MDCK cells, cells were maintained in DMEM supplemented with 10% fetal bovine serum at 37°C and 5% CO_2_. For infection, cells were infected in serum-free infection medium containing TPCK-treated trypsin and incubated at 33°C and 5% CO_2_ for 72 hours. Supernatants were collected, clarified by centrifugation, aliquoted, and stored at −80°C.

### Ferret Infection and Sample Collection

Six-to nine-month-old, female ferrets (*Mustela putorius furo*) were obtained from Triple F farms and acclimated in the UGA animal facilities. All animal experiments were performed in accordance with protocols approved by the University of Georgia Institutional Animal Care Committee (IACUC). For infections, ferrets were anesthetized with isoflurane and intranasally inoculated with 10^6^ PFU of virus diluted in 1mL (500µL per nostril) of sterile PBS. After infection, animals were monitored twice daily for clinical signs. Weights and temperatures were recorded every other day. After each infection, nasal wash samples were collected on days 1, 3, 5, and 7 for analysis of viral replication. While the animals were anesthetized, 3mL of sterile PBS was flushed through their nostrils, collected, and stored at −80°C until further use.

### Plaque Assay

Viral titers for viral stocks and nasal wash samples were quantified using a plaque assay. MDCK cells (FR-926) were seeded in 12-well plates and incubated until they reached 80% confluence. The cells were then inoculated with serial tenfold dilutions of the virus in infection medium (DMEM with 0% FBS) and incubated for 1 hour at 35°C with 5% CO2. Following the infection period, the inoculum was removed, and the cells were overlaid with 2 mL of 0.8% Avicel in MEM supplemented with 1 M HEPES, 200 mM L-glutamine, 7.5% NaHCO3, and 1X antibiotics-antimycotics. The plates were incubated at 35°C with 5% CO2 for 72 hours. After incubation, plaques were visualized by fixing the cells with a methanol:acetone solution (80:20) and staining with crystal violet. Viral titers were determined by counting plaques and were expressed as plaque-forming units (PFU) per milliliter.

### Ferret Cytokine Multiplex Assay

Cytokine expression in nasal wash samples from ferrets was measured using a Luminex multiplex assay kit (Ampersand Biosciences), following the manufacturer’s instructions. In brief, nasal wash samples were diluted 1:2 in assay buffer and added to a 96-well plate along with controls and standards for generating standard curves. Luminex beads conjugated with ferret-specific antibodies and proteins were added to the wells and incubated for 2 hours. Following incubation, a biotinylated detection antibody was added and incubated for 1 hour. Finally, streptavidin-phycoerythrin was added and incubated for 30 minutes. Plates were analyzed using a Luminex MAGPIX system, and cytokine concentrations were determined based on standard curves.

### Multiplex ELISA (human moDCs)

Cytokine and chemokine concentration in the infected and untreated moDCs supernatants was quantified using MILLIPLEX® MAP HUMAN CYTOKINE / CHEMOKINE MAGNETIC BEAD PANEL 96 Well Plate Assay as described by the manufacturer. Cell supernatants were collected post infection at 6h, 24 and 48h time points and stored at −80°C until further analysis. Prior to the assay setup, the supernatants were thawed, vortexed and centrifuged at 1500 rpm for 5 minutes before adding them on microtiter plate provided in the kit. Magnetic capture beads conjugated with analyte-specific antibodies were added to the plate, followed by addition of standards, controls, and samples (duplicates). An 8-point serial dilution of cytokine standards was prepared to generate standard curves. A cytokine magnetic 12-plex panel for the Luminex platform (Millipore Milliplex) was used and consisted of the following analytes: IFN-α, IFN-γ, IL-1b, IL-2, IL-4, IL-6, IL-8, IL-10, IL-12P70, IP-10, MIB-1B and TNF-α. Data acquisition was performed on the Luminex® 200− system which was calibrated and verified prior to each run using calibration and verification beads to ensure assay performance, reproducibility, and batch consistency. xPONENT software was used to visualize and export the acquired data. The cytokine concentrations were then determined using Belysa−Immunoassay Curve Fitting Software V1.1.0 (#40-122). A five-parameter logistic (5-PL) and four-parameter logistic (4-PL) regression model generated by Belysa® was used to visualize the standard curve for each analyte and remove outliers. Post standard curve analysis, the final concentrations were expressed in pg/mL.

### RNA Extraction and Luminex Quantigene Multiplex Assay

Total RNA was extracted from ferret nasal wash samples using the Quick-RNA Miniprep Kit (Zymo Research) following the manufacturer’s protocol. Briefly, samples were lysed in RNA Lysis Buffer, and the lysate was passed through a Zymo-Spin Column to bind RNA. Genomic DNA contamination was minimized by on-column DNase I treatment for 15 minutes at room temperature. The column was then washed with RNA Wash Buffer, and RNA was eluted in RNase-free water. RNA concentration and purity were assessed using a Nanodrop spectrophotometer (Thermo Fisher Scientific. Extracted RNA was stored at −80°C until further use.

Gene expression analysis was performed using the QuantiGene Plex Assay (Thermo Fisher Scientific), a hybridization-based assay that quantifies target RNA without the need for reverse transcription or amplification. Assays were conducted according to the manufacturer’s instructions. Briefly, 200 ng of total RNA per sample was hybridized to target-specific probe sets in a 96-well plate, followed by signal amplification using branched DNA (bDNA) technology. After hybridization and washing steps, target-specific signals were detected using Luminex xMAP technology. Mean fluorescence intensity (MFI) values were normalized to the housekeeping gene glyceraldehyde-3-phosphate dehydrogenase (GAPDH) and used for relative gene expression analysis. The QuantiGene Plex assay was selected for its sensitivity and multiplexing capability, allowing simultaneous quantification of multiple transcripts.

### RNA Isolation and RT-qPCR for Primary Cell Experiments

moDC pellets were collected post infection at 6h, 24h and 48h time points, followed by a cold PBS wash. The cell pellets were lysed using Proteinase K and RNA lysis buffer for 30 minutes at room temperature and stored at −80 °C until the RNA isolation. RNA extraction was performed using Agencourt RNAdvance Cell v2 Total RNA extraction kit (Beckman Coulter), according to the manufacturer’s protocol (Mesnage et al., 2017). The kit uses solid phase reversible immobilization (SPRI) paramagnetic bead technology as indicated in the reference (33). Paramagnetic beads were added to the lysate and bead-RNA complexes were captured using a magnetic stand followed by a series of washes to remove contaminants. The bound RNA was treated with RNase-free DNase I and then eluted using nuclease-free water. The concentration of eluted RNA was determined using Nanodrop 1000 Spectrophotometer V3.8 at 260nm and then stored at −80 °C until the RT-qPCR was performed. RT-qPCR was performed to quantify relative gene expression of Suppressor of Cytokine Signaling 1 (SOCS1) gene in infected and untreated moDCs using the New England BioLabs Luna Universal One-Step RT-qPCR Kit as instructed by the manufacturer. The BioRad 1000C Thermal Cycler was used to perform PCR on the quick plate/SYBR only mode with the following thermocycling reaction: reverse transcription for 1 cycle at 55°C for 10 minutes, initial denaturation for 1 cycle at 95°C for 1 minute, denaturation at 95°C for 10 seconds and extension at 60°C for 30 seconds for 40 cycles. A melt curve step was added to the run reaction to evaluate specificity of amplification. Relative gene expression of SOCS1 was determined by normalizing its Ct values to the housekeeping gene RPS11 and respective mock using ΔΔCT method.

### Pathway Analysis

Differentially expressed genes (DEGs) were analyzed for pathway enrichment using SRplot (https://www.bioinformatics.com.cn). Gene lists were uploaded to the platform, and KEGG (Kyoto Encyclopedia of Genes and Genomes) pathway enrichment analysis was performed using the default parameters. The gene set enrichment analysis (GSEA) function was used to determine statistically significant pathways associated with the DEGs. For GO (Gene Ontology) analysis, genes were categorized into biological process (BP), cellular component (CC), and molecular function (MF) subgroups, and enrichment was assessed using adjusted p-values. Network plots were generated to visualize significantly enriched pathways. Statistical significance was determined using false discovery rate (FDR) correction, and results were interpreted based on adjusted p-values (< 0.05). All visualizations were downloaded directly from SRplot for further analysis and figure preparation.

### Human Primary Immune Cell Isolation and *in vitro* moDC Differentiation

Human primary CD14+ monocytes cells were isolated from buffy coats collected from healthy donors (New York Blood Center) using a standard protocol. Ficoll gradient centrifugation (Histopaque, Sigma-Aldrich) was used to isolate PBMCs from the buffy coats. Followed by this, CD14+ monocytes were isolated from the PBMCs using MACS® Cell Separation CD14+ isolation kit (Miltenyi Biotech) according to the manufacturer’s instructions. Isolated CD14+ monocytes were differentiated in DC media (RPMI medium, 2mM L-glutamine, 100U/mL penicillin, 100 µg/mL streptomycin, 1mM MEM-Sodium pyruvate, 10% (v/v) Hyclonal Fetal Bovine Serum (LifeTechnologies) and was supplemented with 500 U/mL human granulocyte-macrophage colony stimulating factor (GM-CSF) (PeproTech), 1000 U/mL human IL-4 (PeproTech) for 5 days at 37°C and 5% CO2 to obtain monocyte-derived dendritic cell (moDC) phenotype.

### Infection of Primary moDCs

moDCs were mock-infected with infection media (RPMI medium, Penicillin/Streptomycin, 0.2% Bovine albumin serum and TPCK-Trypsin (1µg/mL)) or with virus diluted in infection media to achieve desired multiplicity of infection (MOI). Cells were infected for one hour at 37°C at MOI of 1. After the hour infection, the inoculum was removed and cells were supplemented with appropriate fresh media and incubated at 37°C for up to 24 hours.

### Human PBMC Infection for Cytokine Response and Flow Cytometry

PBMCs were isolated from buffy packs sourced from the Australian Red Cross Lifeblood and whole blood donations from healthy volunteers. These studies were approved by the University of Melbourne (31236, 13344, 1443389). Experiments conformed to the Declaration of Helsinki Principles and the Australian National Health and Medical Research Council Code of Practice.

PBMCs were isolated using density gradient centrifugation, cryopreserved in liquid nitrogen, and stored until further use. For infections, PBMCs were thawed and resuspended in complete RPMI-1640 media supplemented with 10% fetal calf serum (FCS) and enriched supplement. A final number of 1 × 10^6^ cells was added to 24-well plates and infected with an MOI of 1 or 4 in serum-free RPMI-1640 for 1 hr at 37°C with 5% CO_2_. Following incubation, cells were washed to remove unbound virus and resuspended in fresh complete RPMI-1640 media. Infected PBMCs were incubated at 37°C with 5% CO_2_ for either a total of 8 or 22 hrs. After incubation, cells were collected, washed and processed for flow cytometry or gene expression analysis. Mock-infected PBMCs, treated identically but without virus exposure, were used as controls.

### Gene Expression in Human PBMCs

Total RNA was extracted from infected and mock-infected human PBMCs using the RNeasy Mini Kit (Qiagen) according to the manufacturer’s protocol. RNA concentration and purity were determined using a NanoDrop spectrophotometer. Reverse transcription was performed using total RNA with the High-Capacity cDNA Reverse Transcription Kit (Thermo Fisher Scientific), following the manufacturer’s instructions. Quantitative PCR (qPCR) was carried out on a QuantStudio 7 Pro System (Applied Biosystems). Each reaction contained 10 µL of PowerUp SYBR Green Master Mix (Thermo Fisher Scientific), 0.4 µL of 10 µM forward and reverse primers, 2 µL of 5ng/µL cDNA, and 7.2 µL of nuclease-free water. The qPCR conditions were as follows: initial denaturation at 95°C for 2 minutes, followed by 40 cycles of denaturation at 95°C for 15 seconds and annealing/extension at 60°C for 1 minute. Primers targeting IFIT2, IFNγ, SOCS1, MX1, ISG15, IFIT1, IFIT3, TNF, and STAT1 were designed using NCBI Primer-BLAST. GAPDH and 18S rRNA were used as housekeeping genes for normalization. Relative gene expression levels were calculated using as fold change 2^-(ΔΔCt)^, and the geometric mean (GEOmean) of GAPDH and 18S was used as the normalization factor. Data are presented as fold changes relative to mock-infected PBMCs. No-template controls were included to ensure the absence of contamination.

### Flow Cytometry

Peripheral blood mononuclear cells (PBMCs) were prepared as previously described. For viability assessment, cells were first stained with LIVE/DEAD Aqua (LD) viability dye for 15 minutes at room temperature in the dark. Following this, cells were stained for surface markers. Surface antibodies were diluted in MACS buffer (PBS with 5% BSA and 2mM EDTA) and used to target the following markers: CD161 (BV605), CD4 (BV650), CD19 (BV711), CD56 (BV785), CD14 (APC-Cy7), CD8 (PerCP-Cy5.5), TCRvα7.2 (PE), CD3 (PE-CF594), and TCRγδ (PE-Cy7). Staining was performed for 30 minutes on ice in the dark. After incubation, cells were washed with MACS buffer and fixed with BD fixation/permeabilization solution. Next, cells were intracellularly stained with influenza nucleoprotein (NP; FITC) in BD Perm/Wash Buffer for 30 minutes on ice in the dark. Samples were washed with Perm/Wash Buffer and resuspended in MACS buffer, before analyzing using a LSRII Fortessa (BD), and data were processed with appropriate compensation and gating strategies.

For moDC flow cytometry analysis moDCs were collected by fully detaching from the plate via gentle pipetting and transferred to 1.5 mL Eppendorf microcentrifuge tubes. Cells were spun at 1500 G for 5 minutes at RT. Cells were then resuspended in PBS and transferred to a 96-well plate (round bottom) to continue flow cytometry staining. Cells were stained for viability with 100 μL of LIVE/DEAD fixable Blue (1:2000, Thermo) for 15 minutes on ice. Cells were washed with FACS buffer (PBS supplemented with 2% FBS and 5 mM EDTA) and stained with Human TruStain FcX blocking solution (5 μL per 1 x 106 cells) (BioLegend) diluted in FACS buffer for 15 minutes on ice. Cells were then stained for 1 hour on ice with a cocktail of the following antibodies: CD11b (Clone ICRF44, BD Biosciences) and CD209 (9E9A8, BioLegend). After incubation, cells were washed in FACS buffer and centrifuged at 1500 G for 5 minutes. Cells were then fixed with 100 μL of BD Biosciences perm fixation buffer (BD Biosciences) on ice for 20 minutes. After fixation cells were either stained directly with intracellular Influenza B NP Monoclonal Antibody (H89B), FITC (ThermoFisher) or stored at 4 degrees until further staining. Cells were then washed twice in FACS buffer and centrifuged at 1500 G for 5 minutes and resuspended in 200 μL FACS buffer for acquisition on the Cytek Aurora Spectral flow cytometer. Flow cytometry analysis was performed on FlowJo analysis software.

### Statistical Analysis

All statistical analyses were performed using GraphPad Prism version (GraphPad Software, San Diego, CA). Ferret cytokine data were analyzed for statistical differences using a two-way ANOVA. Where no statistical differences were identified, no lines are shown. Gene expression data from human PBMCs were evaluated using an ordinary one-way ANOVA. Flow cytometric data were analyzed using a two-way ANOVA with Geisser-Greenhouse correction. A *P* value of <0.05 was considered statistically significant and is represented as follows: \**P* < 0.05, \*\**P* < 0.01, \*\*\**P* < 0.001, and \*\*\*\**P* < 0.0001.

## Acknowledgements

This work has been funded in whole or in part with Federal funds from the National Institute of Allergy and Infectious Diseases, National Institutes of Health, Department of Health and Human Services under Contract No. 75N93021C00018 S.M.T (NIAID Centers of Excellence for Influenza Research and Response, CEIRR). The work from the Fernandez-Sesma lab was funded by the NIH contract 75N93021C00014 (CRIPT) as part of the CEIRR. This research was funded in whole or part by the National Health and Medical Research Council Investigator Grants: Investigator L2 to K.K. (#2033783).

